# ELITE: Expression deconvoLution using lInear optimizaTion in bulk transcriptomics mixturEs

**DOI:** 10.1101/2023.03.06.531002

**Authors:** Asier Antoranz, Carlos Mackintosh, María Ortiz, Jon Pey

## Abstract

Understanding the cellular composition of tissue samples is crucial for identifying the molecular mechanisms underlying diseases and developing cellular targets for therapeutic interventions. Digital cytometry methods have been developed to predict tissue composition from bulk transcriptomic data, avoiding the high cost associated with single-cell profiling. Here, we present ELITE, a new digital cytometry method that utilizes linear programming to solve the deconvolution problem. ELITE uses as inputs a mixture matrix representing bulk measurements, and a signature matrix representing molecular fingerprints of the cell types to be identified. The signature matrix can be obtained from single-cell datasets or the literature, making ELITE more flexible than other methods that rely solely on single-cell data. We evaluated ELITE on three publicly available single-cell datasets and compared it with five other deconvolution methods, showing superior performance, particularly when there were cell types with similar expression profiles. As a case study, we evaluated the prediction of tumor cellularity using purity estimates from 20 different TCGA carcinoma datasets.

## Introduction

Bulk transcriptomic data represent average expression levels for each gene across different cell types present in a sample, confounding downstream analyses such as differential gene expression or pathway analysis by differences in cell type proportions (Avila Cobos et al., 2020). In the context of cancer, the composition of the tumor microenvironment (TME), a heterogeneous mixture of immune cells, endothelial cells, fibroblasts, and tumoral cells (Denton, A.E., et al., 2018) has shown an active role in tumorigenesis, progression, metastasis, and inflammatory processes triggered by cancer treatments (Hanahan, D., and Weinberg, R.A., 2011). In this line, several studies have shown a positive correlation between tumor infiltrating lymphocytes (TILs) and patient prognosis or response to checkpoint immunotherapy (Sharma, A., et al., 2019; Hendry, S., et al, 2017; Wolf, Y., et al., 2019).

New technologies, including sequencing-based single-cell transcriptomic technologies (e.g., scRNA-Seq), are rapidly evolving allowing cell-type-specific transcriptome profiling (Tang, F., et al., 2009). Nevertheless, they are still in their early stages and, in the case of scRNA-Seq technologies for example, they suffer from low sensitivity and high noise primarily due to a high dropout rate and cell-to-cell variability (Yang, T., et al., 2021). Moreover, the amount of sample needed, and cost associated to single cell sample processing remains relatively high, making the availability of these datasets scarce. This has relevant limitations, particularly when validating novel hypotheses in publicly available datasets where most of the existing data are derived from bulk technologies (RNA-seq and microchip).

Consequently, a myriad of *in-silico* approaches have been developed to infer sample cytometry (cell-type proportions) from bulk transcriptomic data (Abbas, A.R., et al., 2009; Newman, A.M., et al., 2015; Baron, M., et al., 2016; Tsoucas, D., et al., 2019; Wang, X., et al., 2019; Newman, A.M., et al., 2019; Danziger, S.A., et al., 2019; Andersson, A., et al., 2020; Yang, T., et al., 2021). These are often called “digital cytometry” or “deconvolution” methods and try to solve the blind signal separation (BSS) problem. They are typically classified in two categories: regression-based deconvolution algorithms (linear models) and gene set enrichment-based methods (rank-based methods) (Jiménez-Sánchez, A., et al., 2019). Both types of algorithms rely on cell-type-specific markers that are selected according to prior knowledge or derived from the analysis of single-cell data. Deconvolution algorithms use signature matrices containing the average expression of a subset of differentially expressed genes, while enrichment-based methods assign curated gene sets to represent cell types before computing scores as a function of the expression of the genes in each set (Jiménez-Sánchez, A., et al., 2019).

The majority of the methods belong to the deconvolution category and they are normally solved using non-negative least squares (NNLS, Abbas, A.R., et al., 2009), non-negative matrix factorization (NMF, Elosua-Bayes, M., et al., 2021), or support vector regression (SVR, Newman, A.M., et al., 2015) through the maximum likelihood estimation (MLE) or the maximum posterior probability estimation (MAP) to obtain cell fractions (Avila Cobos, F., et al., 2018).

In this work, we present (**E**xpression deconvo**L**ution using l**I**near optimization in bulk transcriptomics mixtur**E**s) ELITE, a new deconvolution method that uses Linear Programming (LP) to solve the BSS problem. ELITE estimates the cell-type fractions of bulk transcriptomic samples using a reference signature matrix that can be obtained from relevant single-cell data, purified cell populations, or predefined signature matrices from the literature (i.e., LM22 (Newman, A.M., et al., 2015), TR4 (Newman, A.M., et al., 2019), TIL10 (Finotello, F., et al., 2019) or others (Aran, D., et al., 2017)). This work focuses exclusively on the deconvolution problem (inferring cell fraction given a signature matrix (B)). Considerations regarding inherent factors such as data normalization and the construction of the signature matrix, specifically marker selection strategies for the genes included in the matrix B, have been extensively reviewed by other authors (Zhong, Y., and Liu, Z., 2012; <u>Hoffmann</u>, M., et al., 2006; Hoggmann, M., et al., 2006; Vallania, F., et al., 2018; Avila Cobos, F., 2020) and fall outside the scope of the presented work. ELITE outperforms competing methods over a range of benchmarked datasets with varying number of cell types.

## Methods

Five subsections have been included to present the methodology behind ELITE:

- *Mathematical model*: mathematical equations modelling the deconvolution algorithm;
- *Datasets*: experimental data used throughout the study;
- *Algorithms*: competing methods included in the study;
- *Internal validation of synthetic bulk samples*: validation experiment performed across three different single-cell datasets;
- *Application to real bulk data*: validation experiment performed in real bulk data by using a publicly available signature matrix.

### Mathematical model

Linear Programming (LP) (Dantzig, G.B., 2002) formulation defines the core of the deconvolution problem. LP is a well stablished modeling technique that has been successfully applied in systems biology for decades (Gusfield, D., 2019). We propose a new mathematical formulation that exploits the inherent potential of the LP to address the bulk gene expression deconvolution problem.

We define a bulk gene expression data matrix (*M*) comprising *G* genes and *S* samples. Let *M_gs_* be the expression value associated with gene *g* (*g = 1,…,G*) in sample s (*s* = *1,…,S*). Similarly, we define the signature matrix (B) comprising G genes and C cell types, being *B_g,c_* the signature coefficient corresponding to gene g and cell type *c* (*c=1,…,C*). In essence, *B_g,c_* reflects the average expression of *g* in cell type *c*. Consequently, and for a given sample *s*, the following set of linear equations applies:

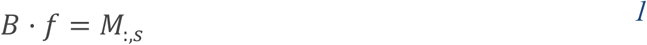

With *f_c_* being the fraction array [f_1_,…,f_c_]. *M_:s_*, is the column vector with the gene expression for sample *s*. From this point onwards, the problem is independently defined and solved for each sample *s*.

Equation 1 provides as many constrains as genes (G):

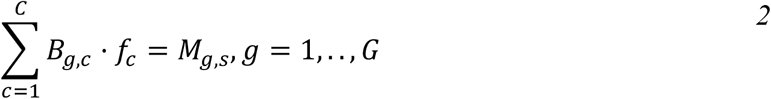

Given the nature of the fraction array, *f_c_*, must include exclusively non-negative values.

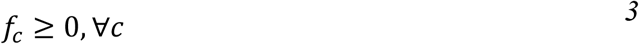

In consequence, the system of equations comprising equations 1–2 results in a convex space where LP has proven to be an extremely powerful tool (Gusfield, D., 2019, Pey, J., & Planes, F. J., 2014). Additionally, this system of linear equations might lead to an infeasibility, where no solution exists that satisfies all the constraints. This is accounted by including an error component, which is addressed in our formulation with two non-negative variables, ε_g_ and δ_g_, which model the negative and positive components of the error respectively. Equation 2, is therefore extended as follows:

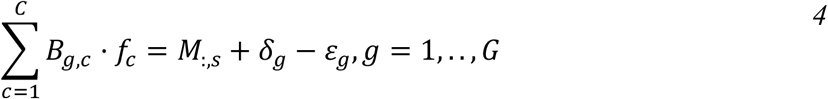

This error component must be accounted for in the objective function when finding a solution.

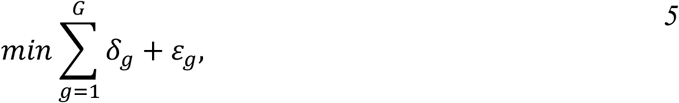

As a result, the LP defined by Equations 3–5, will find the combinations of *f_c_* that minimize the norm-1 of the error slack variables included in Equation 4. Overall, each sample defines a particular LP that is independently solved.

### Datasets

We considered three different single-cell datasets with increasing levels of complexity, one compendium of 20 bulk RNA-seq datasets and one predefined signature matrix in the different analyses included in the present study.

#### Human Pancreatic Islets (PI, Xin, Y., et al., 2016; GSE81608)

This dataset contains 1,492 human pancreatic islet cells divided in four different endocrine cell types (alpha, beta, delta, and gamma) from 18 different samples (6 diabetic and 12 non-diabetic patients).

#### Human Trabecular Meshwork (HTM, Patel, G., et al., 2020)

Contains 8,758 cells classified in 12 different cell types from eight donors. Cell-types include epithelium, melanocytes, smooth muscle cells, macrophages, TNK lymphocytes, three types of endothelial cells (vascular endothelium, lymphatic-like endothelium, and pericytes), 2 types of Schwann cells (Schwann-like and myelinating Schwann cells) and 2 types of TM cells (TM1 fibroblast-like, and TM2 myofibroblast-like). Single cell data obtained from NCBI SRA (accession number PRJNA616025).

#### Mouse lumbar dorsal root ganglion (DRG, Yang, T., et al., 2021)

Includes 3,352 cells from five mice classified in 14 different cell types including endothelial cells, satellite glia, tyrosine hydroxylase-containing neurons (Th), three subtypes of neurofilament containing neurons (NF_Calb1, NF_Pvalb, NF_Ntrk2.Necab2), three subtypes of non-petidergic neurons (NP_Nts, NP_Mrgpra3, NP_Mrgprd), and five subtypes of peptidergic neurons (PEP1_Dcn, PEP1_S100a11.Tagln2, PEP1_Slc7a3.Sstr2, PEP2_Htr3a.Sema5a, PEP3_Trpm8). Data directly obtained from the authors.

#### Tissue Cancer Genome Atlas (TCGA) dataset: carcinomas

These datasets contain bulk gene counts for 8,166 samples from 20 different datasets (Supplementary Table 1). Gene counts were downloaded from the TCGAbiolinks R package (Colaprico, A., et al., 2016) and estimated tumor purity values were obtained from (Aran, D., et al, 2015).

#### TR4 signature matrix

TR4 (Newman, A., et al., 2019) was obtained from FACS-sorted profiles of epithelial cells (EPCAM+), fibroblasts (CD10+), endothelial cells (CD31+) and immune cells (CD45+), obtained from freshly resected surgical tumor samples from patients with Non-Small Cell Lung Carcinoma (NSCLC). It includes the average gene expression of 933 genes for the 4 different cell types. Data was obtained from the publication (Newman, A., et al., 2019).

### Algorithms

ELITE was compared to five other digital cytometry methods including: CibersortX (CSX) (Newman, A.M., et al., 2019), DeconRNASeq (Gong, T., and Szustakowski, J.D., 2013), Non-Negative Least Squares (NNLS, Abbas, A.R., et al., 2009), MuSiC (Wang, X., et al., 2019) and AdRoit (Yang, T., et al., 2021). While ELITE, CSX, DeconRNASeq and NNLS use a signature matrix with averaged gene expressions per cell type for the deconvolution problem, MuSiC and AdRoit directly exploit single-cell data for the construction of their respective models. Selection of these methods for benchmarking was based on diversity of their mathematical approaches.

#### CibersortX (CSX)

(Newman, A.M., et al., 2019) uses a v-support vector regression, a machine learning technique robust to noise, with unknown mixture content and collinearity among cell type reference profiles (Newman, A.M., et al., 2015). CSX implementation is coupled with a batch correction scheme to adjust for cross-platform differences (CSX B-mode).

#### DeconRNASeq

(Gong, T., and Szustakowski, J.D., 2013) solves a non-negative least-squares constrain with quadratic programming to obtain the global optimal solution to estimate the mixing fractions of distinctive cell types in bulk samples. DeconRNASeq R package (version 1.40.0) was downloaded from Bioconductor.

#### Non-Negative Least Squares (NNLS)

(Abbas, A.R., et al., 2009) applies a linear least squares algorithm followed by removal of the lowest negative coefficients from the equation and iterates until all coefficients are non-negative. This method was implemented with the MuSiC R package (v1.0.0) obtained from GitHub (https://github.com/xuranw/MuSiC).

#### MuSiC

(Wang, X., et al., 2019) uses a gene weighting strategy which prioritizes consistent genes across subjects up-weighing genes with low cross-subject variance and downweighing genes with high cross-subject variance. This makes unnecessary the pre-selection of marker genes allowing for larger numbers of genes to be considered in the deconvolution. Weights are coupled with weighted NNLS to predict cell fractions in the mixed sample. MuSiC R package (v1.0.0) was obtained from GitHub (https://github.com/xuranw/MuSiC).

#### AdRoit

(Yang, T., et al., 2021) uses an adaptative learning approach to estimate gene-wise corrections, alleviating the platform-related batch differences between the single-cell and the bulk expression data, enhancing cross-platform readout comparability. It also incorporates regularization to reduce collinearity among closely related cell subtypes. To the best of our knowledge, AdRoit is not publicly available (https://github.com/TaoYang-dev/AdRoit). Therefore, we extracted the outputs produced by AdRoit in the comparisons directly from the supplementary data of the original publication.

### Internal validation of synthetic bulk samples

We validated ELITE using three different single-cell datasets with increasing levels of complexity (rising numbers of more similar cell types). Each dataset was analyzed independently but all of them followed the same analytical pipeline which included: (i) single-cell data normalization and quality control, (ii) construction of signature matrices, and (iii) recovery of cell fractions in synthetic bulk samples (pseudobulks). Quality control, construction of signature matrix and recovery of cell fractions in pseudobulk samples was performed following a leave-one-out cross-validation (LOOCV) scheme. That is, when estimating the fractions of a given sample, this sample was excluded from the signature matrix construction.

#### Single-cell data normalization and quality control

First, systematic differences between samples of the same dataset were removed. For that, top 2000 variable features were selected for each sample using Seurat’s FindVariableFeatures function (vst algorithm). Then, top 2000 features repeatedly variable across datasets were picked using the SelectIntegrationFeatures function (standard parameters). Integration anchors were estimated using the FindIntegrationAnchors function (standard parameters). Finally, data from different samples were integrated using the IntegrateData function (standard parameters). After integration, cells with negative count values were trimmed to 0.

For quality control we followed the pipeline described in (Avila Cobos, F., et al., 2020) with some adaptations. Briefly, we removed all samples with less than 30 cells from the analysis. We also removed genes with zero variance. Finally, we kept only genes with a read count greater than 0 for at least 75% of the cells of at least 1 cell type. These subsets of genes represented the candidates for the construction of the signature matrices.

#### Construction of signature matrices

In this passage, we also slightly adapted the methodology described in (Avila Cobos, F., et al., 2020). Briefly, for each gene, the 2 cell types with the highest average expression were calculated (rank1 and rank2). Using these two cell types, we estimated the logFC, the adjusted p-value using the Benjamini Hochberg adjustment method and the area under the ROC curve (AUC). We applied the following cutoffs for each parameter: logFC > 0.25, pAdj < 0.05, AUC > 0.7. From the subset of selected genes, we evaluated if all the cell types were represented in the rank1 group. For those that were not, we included the top 10 genes with the highest relative average value of the non-represented cell types. The signature matrix was then calculated as the average expression of each selected gene in each cell type.

#### Recovery of cell fractions in synthetic bulk samples

Finally, the synthesized bulk data were constructed by summating the expression of all the cells for each gene of the testing sample. We evaluated the recovery of known cell fractions of each method using five complementary statistics: Pearson and Spearman correlation coefficients (PCC and SCC), mean absolute error (MAE), and the root mean square error (RMSE). Additionally, we calculated the norm-2 differences between the input pseudobulk (M) and the recovered pseudobulk (M’=B*f). The first four metrics are based on the differences between the estimated fractions (f) and the real fractions (f’). While PCC represents the linear correlation between the estimated results and the ground truth, SCC indicates the correlation in the ranking even if the relationship is not linear (monotonic relationship). SCC avoids the artificial high correlation scores driven by outliers when the majority of the cell fractions are relatively small (Yang, T., et al., 2021). Thus, both correlation statistics are complementary. The MAE and RMSE represent the norm-1 and norm-2 error. While the norm-1 is more resistant to outliers and therefore more robust, the norm-2 is more resistant to horizontal adjustments and therefore more stable. The fifth metric (norm-2) is a purely mathematic statistic representing the fraction of bulk sample (M) that we can explain with the deconvolution (M’=B*f).

### Application to real bulk data

We validated ELITE in real bulk samples using a compendium of 8,166 cancer samples from 20 datasets of different types of carcinomas from the Tissue Cancer Genome Atlas (Supplementary Table 1). As signature matrix, we used the publicly available TR4 signature (Newman, A.M., et al., 2019) which includes the averaged gene expression of four distinct populations: stromal (CD10), endothelial (CD31), immune (CD45), and tumoral (EPCAM). The analysis is focused on the recovery of fraction of tumoral cells by leveraging tumor-purity annotations obtained from (Aran, D., et al, 2015). Since the mixture and signature matrices were constructed using different technologies, there might exist a batch effect that affects the quality of the deconvoluted fractions. To address this issue, we have developed a multi-step process that goes as follows (Figure 1):

i. **Obtain the initial cell fractions** (f_0_) by applying ELITE on the raw bulk data (M_0_) using the raw signature matrix (TR4_0_).
ii. **Generate synthetic mixture samples** (*M*_*pseudo*_0__) by multiplying N (N=100) vectors representative of the obtained fractions (f’_0_) in the real population times the columns of the gene signature matrix TR4_0_ (M_pseudo0_ = TR4_0_ · f’_0_). To make the gene expression of each pseudobulk sample different, the representative fractions (f’_0_) were obtained as follows:

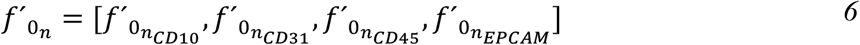 Where *f’*_0*n*__*CD*10_ is a random value sampled from a normal distribution with average equal to the mean fraction of CD10 in f_0_ and standard deviation (sd) equal to the sd of CD10 in f_0_.
iii. **Correct platform-related differences** between M_0_ and the combination of M_pseudo0_ and TR4_0_ (*M*_*combined*0_]) using ComBat-seq (Zhang, Y., et al., 2020) as implemented in the sva R package (Leek J.T., et al., 2022).
iv. **Apply ELITE to the batch corrected bulk samples** (M_1_) using the batch corrected TR4 (TR4_1_) and obtain new fractions (f_1_),
v. **Repeat the process N times** (N=100) using each time the fractions obtained from the previous iteration (f_n-1_, the raw bulk expression matrix M_0_, and the raw signature matrix TR4_0_ to generate a new *M_pseud__On_*, TR4_n_, and M_n_, matrix.
vi. **Select the optimal solution** using the elbow criterion on the estimated cell fractions. To that end, first we estimate the average fraction of each cell type across all samples in each iteration, then, we invert the populations having an increasing trend aligning the direction of all the populations. Fractions and iterations are normalized (range 01) to be on the same units and range. Finally, the normalized values across all cell types were summarized per iteration and the elbow criterion applied to select the optimal iteration (iteration with the closest distance to the (0,0) point).

**Figure 1.**
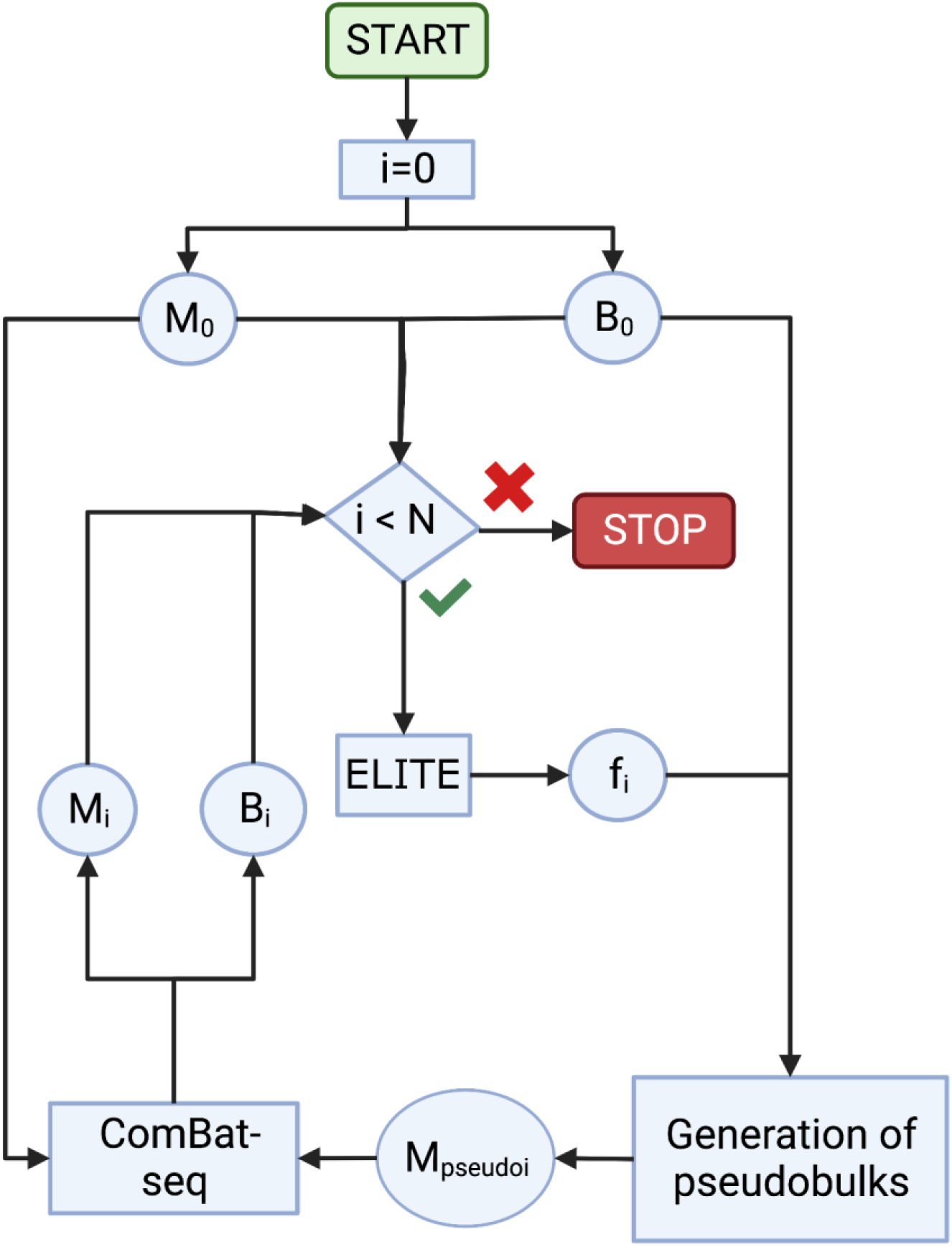
Validation scheme of real bulk samples. Flow diagram showing the scheme followed for the validation of the bulk data. Starting from a specific experimental picture (mixture matrix M_0_ and signature matrix B_0_), we calculate the cell fractions with ELITE (f_i_, here i = 0). Then, we calculate the pseudobulk mixture matrix (M_pseudo_i__) by multiplying 100 vectors representative of the obtained fractions f’i times the columns of the signature matrix. Finally, we merge the original signature matrix B0 and M_pseudo_i__. and apply ComBat-seq between M0 and the merged matrix obtaining Mi and Bi (here i = 1) that will be used to calculate again fi with ELITE. We repeat this process N times (here N = 100) and select the optimal solution by applying the elbow criterion.

Predicted EPCAM fractions in the optimal iteration were correlated with tumor purity estimates provided by (Aran, D., et al, 2015), which includes tumor purity estimates from 5 different methods: ESTIMATE, based on gene expression profiles of 141 immune genes and 141 stromal genes (Yoshihara, K., et al., 2013); ABSOLUTE, which uses somatic copy-number data (Carter, S.L., et al., 2012); LUMP (leukocytes unmethylation for purity), which averages 44 non-methylated immune-specific CpG sites (Aran, D., et al, 2015); immunohistochemistry (IHC) as estimated by image analysis of haematoxylin and eosin stain slides produced by the Nationwide Children’s Hospital Bioscpecimen Core Resource (Aran, D., et al, 2015); and Consensus measurement of Purity Estimations (CPE), which is the median purity level after normalizing levels from all methods to give them equal means and standard deviations (Aran, D., et al, 2015). ELITE’s performance at baseline and optimal iteration were compared with that of CSX (with and without B-mode) by means of the same performance metrics that were described for in the previous section (PCC, SCC, MAD, RMSE, norm-2).

## Results

In this section we show the results obtained with the proposed methodology in two different scenarios: (i) recovering cell fractions in synthetic bulks derived from pooling cells from single-cell datasets and (ii) recovering the tumor fraction of real bulk samples. For each of these scenarios, we compare the performance with state-of-the-art methods. In the last part of the results, referred to as ‘Evaluating corrected Signature Matrices for Carcinomas’, the biological relevance of the modifications in the signature matrix introduced by the suggested batch effect correction is evaluated.

### Recovery of cell fractions in synthetic bulk samples

We compared ELITE to CibersortX (CSX), DeconRNAseq, NNLS, MuSiC, and AdRoit using three different single-cell datasets of increasing complexity (PI, Xin, Y., et al., 2016; HTM, Patel, G., et al., 2020; DRG, Yang, T., et al., 2021) and a Leave-One-Out Cross-Validation (LOOCV) scheme. A summary of the results is reported in Figure 2.

**Figure 2.**
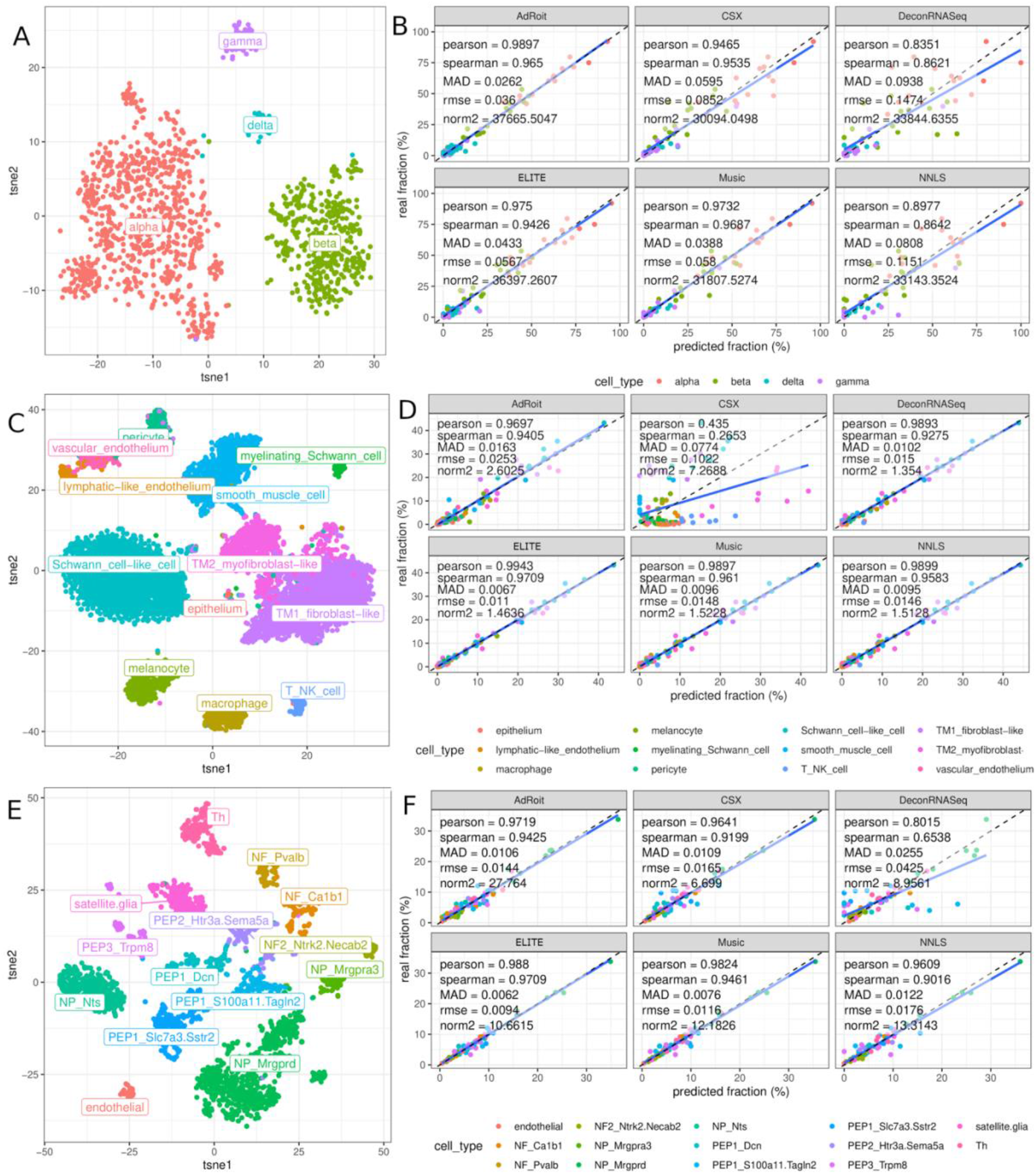
Recovery of cell fractions in synthetic bulk samples. Dimensionality reduction plots (tsne, left column) and cytometric recovery (scatter plot, right column) of pseudobulk samples generated with 3 different datasets: human pancreatic islets (PI, Xin Y., et al., 2016) (A, B), human trabecular meshwork (HTM, Patel, G., et al., 2020) (C, D) and mouse lumbar dorsal root ganglion (DRG, Yang, T., et al., 2021) (E, F).

We started by applying ELITE to a simple human pancreatic islet dataset (Xin, Y., et al., 2016) containing 1,492 cells derived from 18 donors and classified in 4 different cell types (Alpha, Beta, Delta, Gamma). 3 samples (“Non T2D 2”, “Non T2D 7”, “Non T2D 4”) were removed due to low cell count (24, 27, and 27 respectively). From the initial 30,371 genes included, only 1,929 passed quality control (QC). These were then used to generate the t-SNE plot (Figure 2A) and considered as candidates for the construction of the signature matrix B. Pseudobulk samples (matrix M) were constructed by summating the expression of all the cells for each gene in each sample. A leave-one-out cross-validation (LOOCV) scheme was applied, where the fractions for sample *i* were calculated using a signature matrix (B) estimated with *S* \ {*i*} (all the samples from dataset S except for *i*). In this dataset AdRoit had the best performance for the PCC (=0.9897), MAD (=0.0262), and RMSE (=0.036), while MuSiC had the best SCC (=0.9674) and CSX the best norm-2 error (=30,092.38). ELITE, AdRoit, and MuSiC all had very comparable results with CSX positioned slightly worse, and DeconRNASeq and NNLS performing significantly worse than the rest (Figure 2B).

We then evaluated the performance of ELITE in a more complex human trabecular meshwork (HTM) dataset (Patel, G., et al., 2020) containing 8,758 cells derived from 8 donors and classified in 12 different cell types (epithelium, lymphatic-like endothelium, macrophage, melanocyte, myelinating Schwann cell, pericyte, Schwann cell-like cell, smooth muscle cell, T NK cell, TM1 fibroblast-like, TM2 myofibroblast-like and vascular endothelium). From the initial 17,757 genes included in the dataset 1,725 passed quality control and were used to generate the t-SNE plot represented in Figure 2C and considered as candidates for the construction of the signature matrix B. Sample fraction recovery was evaluated using the same strategy defined above (pseudobulk construction + LOOCV). The complexity of this dataset is due to the existence of 3 similar types of endothelial cells, 2 types of TM cells, and 2 types of Schwann cells. In this case, ELITE outperformed all the competing methods in all metrics (PCC=0.9943, SCC=0.9709, MAD=0.0067, rmse=0.011) apart from norm-2 error where DeconRNASeq performed better (=1.354) (Figure 2D).

We also evaluated ELITE in a scenario with multiple similar cell types in a complex mouse lumbar dorsal root ganglion (DRG) dataset (Yang, T., et al., 2021) containing 3,352 cells derived from 5 mice and classified in 14 different cell types (endothelial, NF Ca1b1, NF Pvalb, NF2 Ntrk2.Necab2, NP Mrgpra3, NP Mrgprd, NP Nts, PEP1 Dcn, PEP1 S100a11.Tagln2, PEP1 Slc7a3.Sstr2, PEP2 Htr3a.Sema5a, PEP3 Trpm8, satellite glia, and Th). From the initial 17,271 genes included in the dataset, 1,714 passed QC and were used to generate the t-SNE plot represented in Figure 2E and considered as candidates for the construction of the signature matrix B. Sample fraction recovery was evaluated using the same strategy defined above (pseudobulk construction + LOOCV). The complexity of this dataset derives from the existence of several types of neurofilaments containing neurons (NF subtypes), three subtypes of non-peptidergic neurons (NP subtypes), and five subtypes of petidergic neurons (PEP subtypes). In this case, ELITE outperformed all the competing methods in all metrics (PCC=0.988, SCC=0.9709, MAD=0.0062, rmse=0.0094, norm-2=6509.99) (Figure 2F).

### Recovery of tumor purity in real bulk samples

Finally, we evaluated the ability of ELITE to infer cellular proportions in real bulk samples. We applied ELITE to 8,166 TCGA samples from 20 different carcinoma datasets (Supplementary Table 1).

To estimate the fraction of tumoral cells we used the publicly available TR4 signature matrix (Newman, A.M., et al., 2019) which includes the four main populations found in tumoral samples: stromal (CD10), endothelial (CD31), immune (CD45), and tumoral (EPCAM). We set the focus on carcinomas due to the origin of EPCAM. Therefore, gliomas (GBM, LGG), melanomas (SKCM, UVM), or lymphomas (DLBCL) were excluded from the analysis.

As no gold-standard measures were available for these samples, we considered previous estimates of tumor purity (Aran, D., et al, 2015) available for a subset of 6,755 TCGA samples from 16 carcinomas that were used to further assess the validity of ELITE’s results. 3,611 samples had available purity information from ABSOLUTE, 6,707 from CPE, 5,927 from ESTIMATE, 6,725 from IHC, and 4,907 from LUMP. These purity estimations represent an imperfect gold standard with significant discrepancies across the different methods (Supplementary Figure 1). Due to the low correlation between IHC and the rest of purity estimation methods, IHC was not considered for downstream analysis (Supplementary Figure 1).

Since the signature matrix (TR4) and bulk measurements were obtained using different platforms, we applied the batch-correction scheme described in the methods (Figure 1). Briefly, we calculated the initial cell fractions using the raw uncorrected data, reconstructed a pseudobulk matrix by multiplying the estimated fractions times the signature matrix, used ComBat-seq to correct for batch effect and used the corrected data to estimate new fractions. We repeated the process 100 times and used the elbow criterion to select the optimal iteration (convergence of predicted fractions). The cell fraction evolution graphs and optimal iterations are represented in Supplementary Figure 2. Supplementary Figure 3 shows how this approach nicely reduced the batch-effect and integrated the data.

Initially, we compared the predicted EPCAM fractions with the tumor purity estimates for each iteration and tumor purity estimation method independently. We evaluated the performance using the same four statistics used for the validation of the single-cell datasets (PCC, SCC, MAD, and RMSE). Regardless of the purity estimation method, the optimal iteration returned by the unsupervised criterion defined above (Supplementary Figure 2) aligned with the optimal iteration when evaluating these metrics (Supplementary Figure 4). Additionally, some tumor purity estimation methods aligned much more consistently with the predictions provided by ELITE than others regardless of the iteration. In the optimal iteration, ESTIMATE had the highest correlation values (PCC_ESTIMATE_=0.718, SCC_ESTIMATE_=0.755) and the highest error metrics (MAD_ESTIMATE_=23.2, RMSE_ESTIMATE_=30.1) while ABSOLUTE had the lowest error metrics (MAD_ABSOLUTE_=18.8, RMSE_ABSOLUTE_=23.9) but also lower correlation values (PCC_ABSOLUTE_=0.536, SCC_ABSOLUTE_=0.543). LUMP had comparable error rates to ESTIMATE (MAD_LUMP_=23, RMSE_LUMP_=28.9) but lower correlation values (PCC_LUMP_=0.4, SCC_LUMP_=0.392). Finally, CPE, as expected, had intermediate error rates (MAD_CPE_=20.1, RMSE_CPE_=26.3) and intermediate correlation values (PCC_CPE_=0.601, SCC_CPE_=0.605) since its values are calculated by averaging the purity values of the previous methods (Figure 3A).

**Figure 3.**
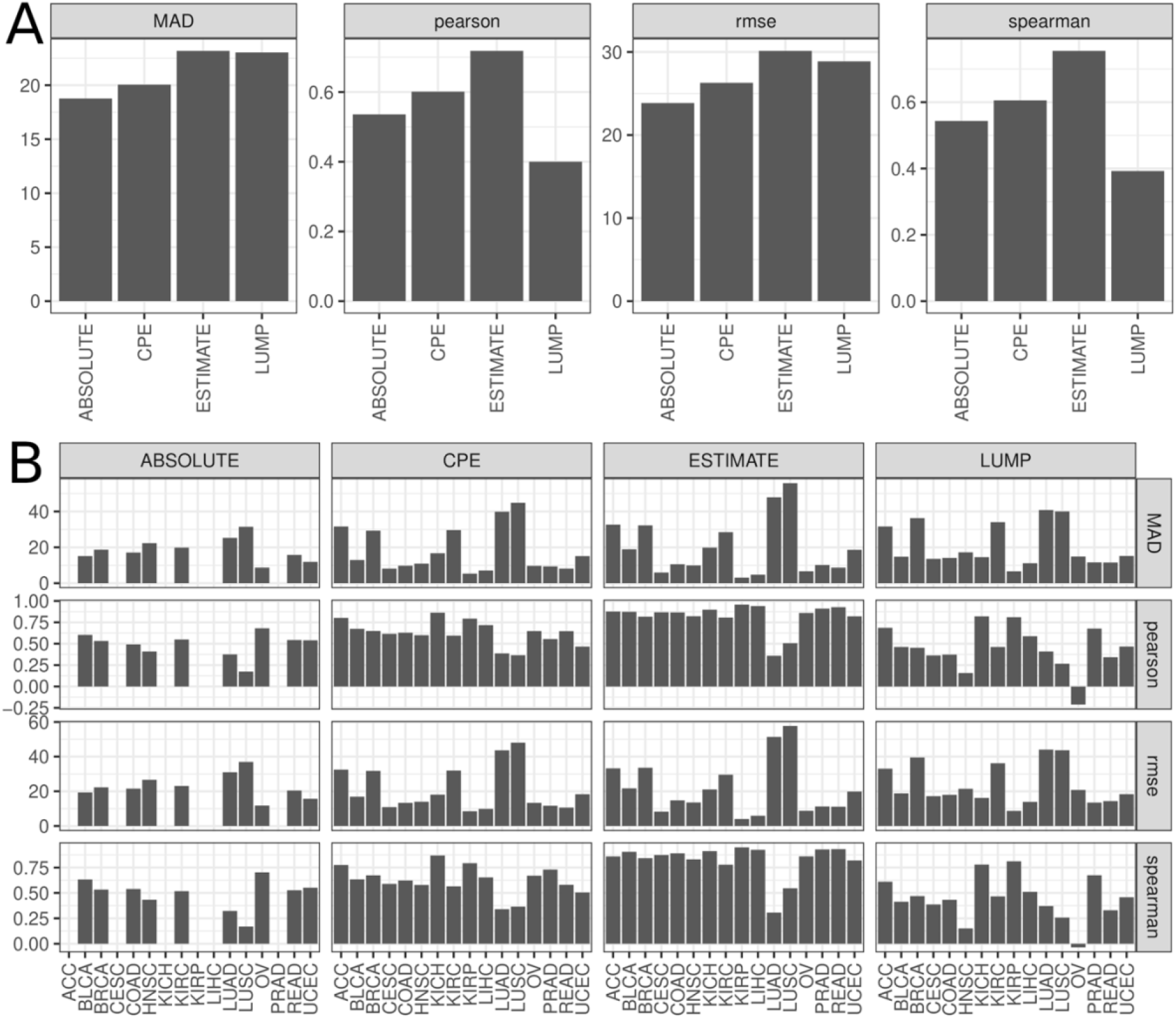
Recovery of real bulk samples using different purity estimation methods in optimized ELITE. A) Barplots showing (from left to right) Median Absolute Error (MAD), Pearson Correlation Coefficient (pearson), Root-Mean-Square Error (RMSE), and Spearman Correlation Coefficient (spearman) for the different tumor purity estimation methods (ABSOLUTE, CPE, ESTIMATE, and LUMP) by pulling together all the samples from the 20 different carcinoma datasets. B) Barplots showing the same results after segmenting by dataset.

When segmenting by dataset, significant differences arise (Figure 3B). To assess which purity estimation method aligned better with ELITE’S results we calculated the average ranking of each metric in each dataset and summarized the results. The method aligning best was ESTIMATE (average ranking = 1.77) followed by CPE (average ranking = 2.08), ABSOLUTE (average ranking = 2.55) and finally LUMP (average ranking = 3.06). Regardless of the purity estimating method, the two worst performing datasets were LUAD and LUSC with the lowest correlation values (in ESTIMATE, PCC_LUAD_=0.359, PCC_LUSC_=0.506, SCC_LUAD_=0.307, SCC_LUSC_=0.546) and highest error rates (in ESTIMATE, MAD_LUAD_=47.9, MAD_LUSC_=55.8, RMSE_LUAD_=51.4, RMSE_LUSC_=57.5). On the other hand, the two datasets with the best recovery were KIRP (in ESTIMATE, PCC_KIRP_=0.958, SCC_KIRP_=0.949, MAD_KIRP_=3.11, RMSE_KIRP_=4.05) and LIHC (in ESTIMATE, PCC_LIHC_=0.0.94, SCC_LIHC_=0.925, MAD_LIHC_=4.74, RMSE_LIHC_=5.93).

We also compared the recovery of ELITE in the raw and optimal iteration with that obtained with CSX. Specifically, we implemented CSX using two different configurations: B-mode (batch correction), and no-B mode (without batch correction). Overall, optimized ELITE had the best performance (average rank = 1.82, Wilcoxon rank-sum test vs CSX no-B-mode (FDR corrected): p-value = 3.57e-09, vs CSX B-mode: p-value = 1.246e-07, vs raw ELITE: p-value = 2.16e-04) followed by raw ELITE (average rank = 2.19, Wilcoxon rank-sum test vs CSX no-B-mode (FDR corrected): p-value = 2.712e-14, vs B-mode CSX: p-value = 5.235e-06), B-mode CSX (average rank = 2.83, Wilcoxon rank-sum test vs CSX no-B-mode (FDR corrected): p-value = 0.004), and lastly no-B-mode CSX (average rank = 3.16). Both methods improved their recovery when considering platform-related differences. However, the improvement introduced by ELITE was significantly more notorious highlighting the success of the suggested batch correction scheme described in Figure 1. In the following lines we describe the results for MAD using ESTIMATE as purity estimation method (Figure 4A). The comparison between the different deconvolution approaches for all the statistics and all the purity estimation methods can be found in Supplementary Figure 5. In 13 out of the 16 TCGA carcinoma datasets where we had purity estimation optimized ELITE provided the best result. Only in LUAD, LUSC, and UCEC the best performance was obtained by CSX b-mode (batch corrected). As previously mentioned, the two best performing datasets were KIRP (MAD_ELITEopt_=3.11, MAD_ELITEraw_=19.3, MAD_CSXbmode_=11.6, MAD_CSXnoBmode_=14.4) and LIHC (MAD_ELITEopt_=4.74, MAD_ELITEraw_=28.5, MAD_CSXbmode_=32.6, MAD_CSXnoBmode_=30) where the batch correction introduced a reduction of 83.9% and 83.4% of the MAD respectively.

**Figure 4.**
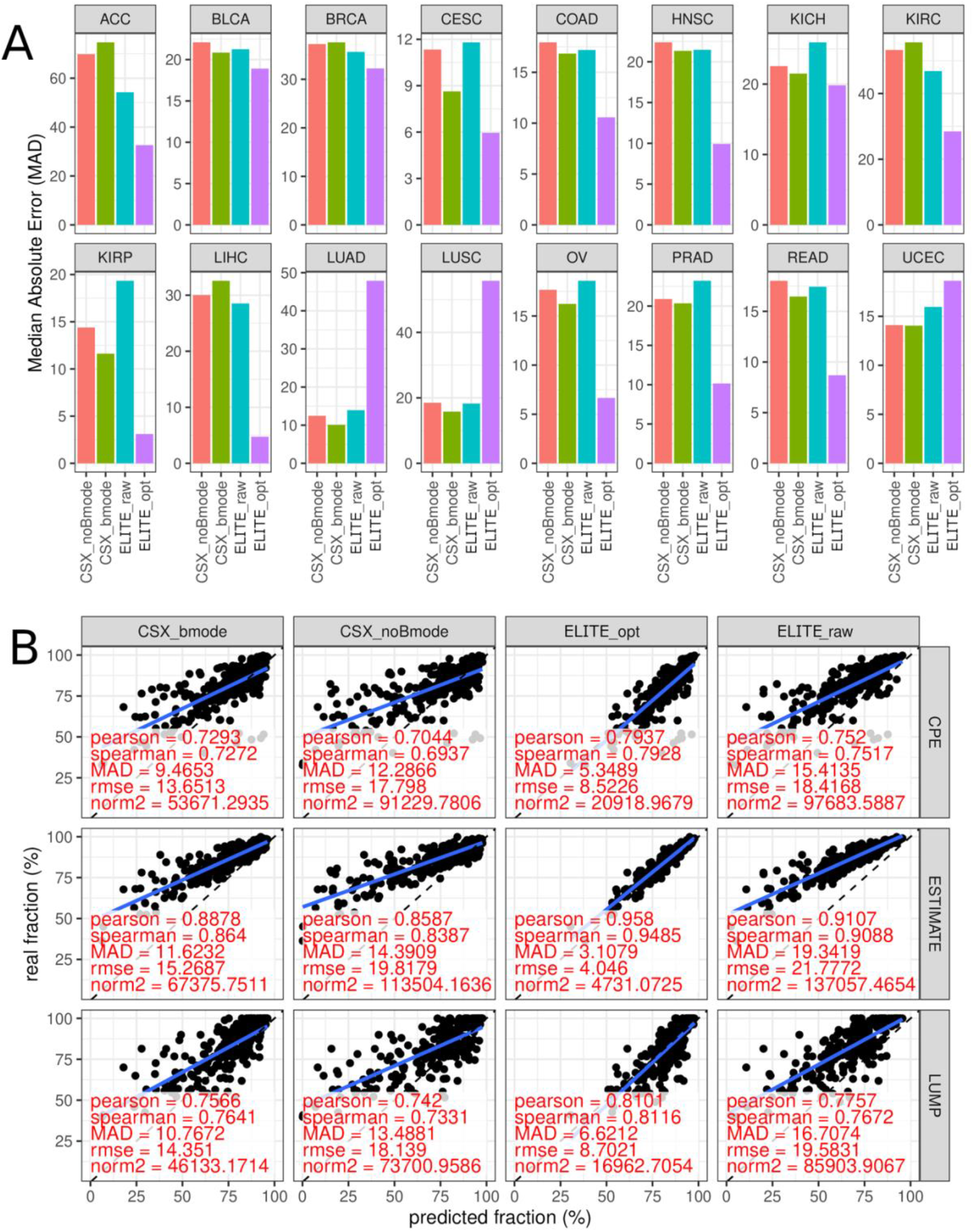
Comparison between different deconvolution methods. A) Barplot showing the Median Absolute Error (MAD) comparing the purity estimations of ESTIMATE with the tumor fractions of CSX without batch correction (CSX_noBmode), CSX with batch correction (CSX_bmode), ELITE without batch correction (ELITE_raw) and ELITE with batch correction (ELITE_opt). B) Scatter plot showing the predicted fraction of tumoral cells (x-axis) versus the estimated tumor purity (y-axis) for different tumor purity estimation methods (CPE, ESTIMATE, LUMP) and different deconvolution methods (CSX_noBmode, CSX_bmode, ELITE_raw, ELITE_opt) for the KIRP dataset. In this dataset, the estimations from ABSOLUTE were not available in Aran, D., et al., 2015.

### Evaluating corrected Signature Matrices for Carcinomas

In the previous section, we applied ELITE to 20 different carcinomas from TCGA (https://www.cancer.gov/tcga). The batch removal pipeline described in Figure 1 was executed obtaining a set of corrected and particularized TR4 carcinoma signatures. Results showed that the application of a batch effect removal process improves purity predictions for both deconvolution methods (CSX and ELITE). Below, we evaluate the biological impact of the correction pipeline implemented in ELITE.

TR4 (Newman, A.M., et al., 2019) was considered as initial template signature matrix. However, the application of ELITE together with the batch effect correction, generated carcinoma-specific TR4 signature matrices with different gene profiles for each tumor type. To assess the degree of variation introduced by the proposed batch-correction framework, the most expressed genes were selected for each cell type and the correlation between the values in the raw and corrected matrices were compared. Unsurprisingly, the signature of the tumoral cells (EPCAM) was the most affected one (PCC = 0.38) as opposed to all the other cell types: CD10 (PCC = 0.9), CD31 (PCC = 0.78), and CD45 (PCC = 1) (Figure 5A). If we split these correlations by dataset (Figure 5B), there is a high variability between the correlation values in the different datasets. For example, LUAD and LUSC, the two carcinoma types that were used to create the TR4 matrix have significantly higher correlation values in the gene expression profile of EPCAM (PCC = 0.84 and PCC = 0.85 respectively) than the rest of datasets (average PCC = 0.388).

**Figure 5.**
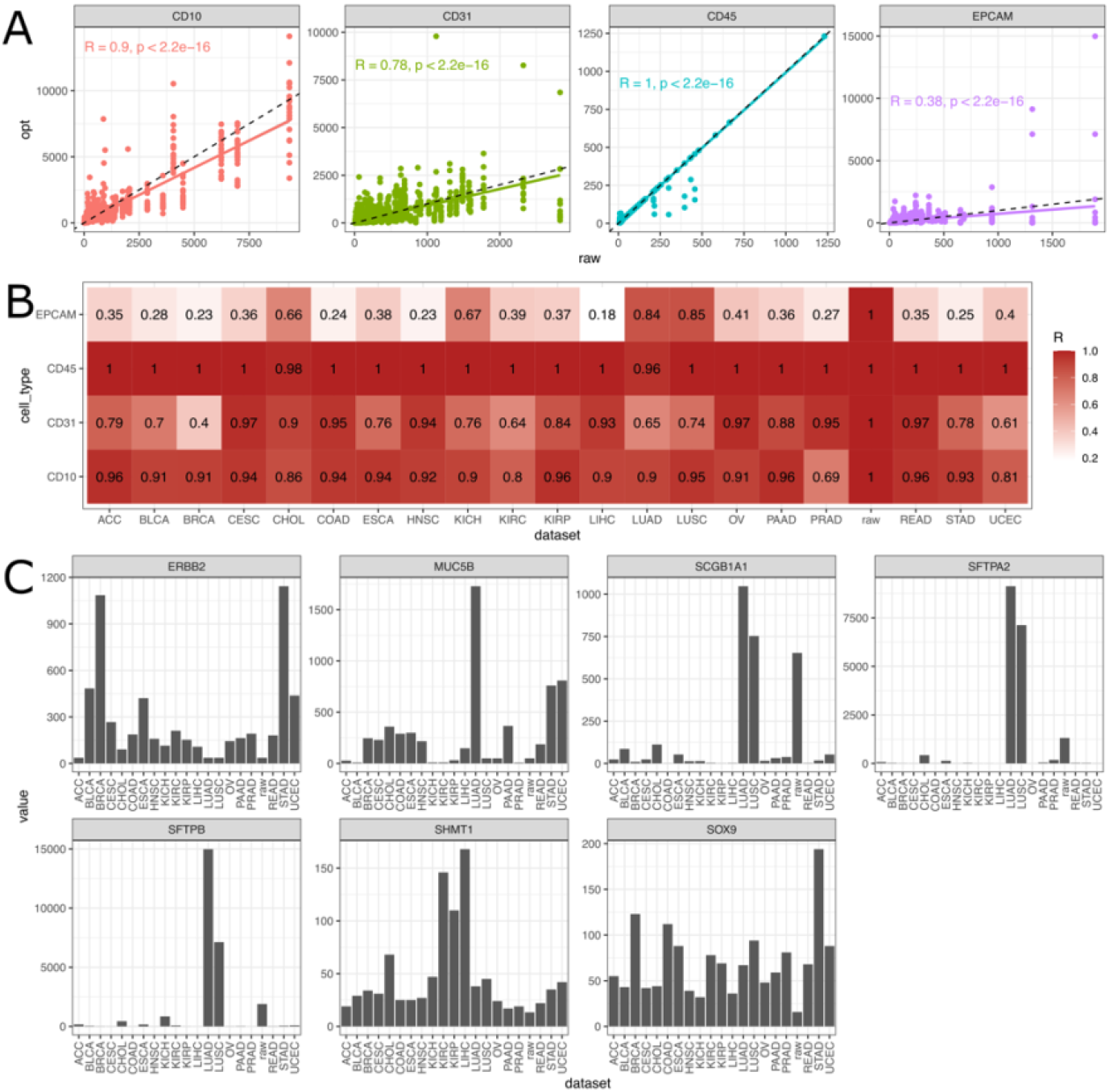
Evaluation of changes introduced by the proposed batch correction framework in the signature matrix. A) Scatter plot showing the expression value in the raw (x-axis) and optimized (y-axis) signature matrix following the batch-effect correction scheme described in Figure 1. B) Heatmap showing the Pearson correlation value between gene expression values in the raw and optimized TR4 segmented by dataset. C) Bar plot showing the expression of the most variable genes in the tumoral profile (EPCAM) for the raw and the carcinoma-specific TR4 signature matrices.

Then, we wanted to understand if the modifications introduced by the method were also relevant from a biological point of view. Table shows the top 10 genes and carcinomas leading to the most significant deviations from the original TR4 tumor signature. Table A and Table B show the top 10 results, sorted by log fold change, whose coefficients in the tumoral signature of TR4 were reduced and increased respectively. Figure 5C presents the expression of these genes for TR4 in the raw and carcinoma-optimized signature matrices for the EPCAM column.

Figure 5C shows how SCGB1A1, SFTPA2, and SFTPB have significantly higher values in LUAD and LUSC, moderate values in the raw TR4 and neglectable values in the remaining datasets (also validated by data in https://www.proteinatlas.org). Based on the hypothesis leading to the definition of the TR4 signature matrix, we expected lung-specific genes, i.e., genes whose expression is high for NSCLC but not for the carcinomas in the corresponding row of Table A:

- **SFTPA2:** Surfactant protein A2. Genetic defects in SFTPA2 have been associated with pulmonary fibrosis and lung cancer (Wang, Y., et al., 2009).
- **SFTPB**: Surfactant Protein B. It has been shown to have diagnostic biomarker ability for lung cancer prediction (Sin, D.D., et al., 2013).
- **SCGB1A1**: Secretoglobin family 1A member 1. Its expression has been associated with alveolar inflammation and immunity (Xu, M., et al., 2020).

Considering the genes included in Table B, the top 10 gene/carcinoma pairs that most increased their expression after correction include 4 different genes (MUC5B, ERBB2, SHMT1, and SOX9). Figure 5C shows how the expression of these genes is not as high for LUSC and LUAD as opposed to other carcinomas. For instance:

- **MUC5B** turns out to be relevant for LUAD. This may seem counterintuitive, as LUAD, together with LUSC samples, composed the samples to reconstruct TR4. However, this gene is highly expressed in LUAD when compared to LUSC (Zhan, C., et al., 2015) and, precisely, this is properly captured by the proposed batch correction pipeline as shown in Figure 5C. In addition to LUAD, MUC5B also results relevant for UCEC and STAD. Previous works support this observation for both carcinomas too, namely UCEC (Hebbar, V., et al., 2005) and STAD (Pinto-de-Sousa, J., et al., 2004).
- **ERBB2** is relevant for several carcinomas, being of especial importance in Breast Carcinoma (Harari, D., et al., 2000). However, evidences are found that ERBB2 is overexpressed in, at least a significant proportion of the patients, for the other tumors in Table B as well: STAD (Albarello, L., et al., 2011) – BLCA (Kaufman, D.S., et al., 2009) - UCEC (Diver, E.J., et al., 2015) and ESCA (Albarello, L., et al., 2011). The Human Protein Atlas (Sjöstedt, E., et al., 2020) (tissue section) provides a graphic landscape of the expression of this gene across different tumors, showing that respiratory carcinomas show a lower expression of ERBB2 than a significant portion of tumors included in the analysis. This supports the need to increase the relevance of ERBB2 in the signature matrix of several carcinomas.
- **SHMT1:** This gene plays a well-known role in the biosynthesis of glycine from serine, which mainly occurs in liver and kidney (Simmons, R.M., et al., 2020), supporting the results included in Table B.

Overall, the unsupervised procedure in Figure 1 successfully captures the following scenarios:

- The relative importance corresponding to lung specific genes is reduced (genes included in Table A).
- The global relevance of genes that are highly specific of a carcinoma other than NSCLC is increased in the particularized TR4 matrix, *e.g*., ERBB2 or SHMT1 in Table B.
- Finally, even though TR4 was generated from NSCLC samples, some genes capture a significant variability across the different tumor subtypes and lead to a correction of the weights in the specific signature matrix (Gene MUC5B in Table B).

These modifications induce a better performance when recovering tumor purity from bulk RNAseq samples in a variety of carcinoma datasets (Figure 4A).

## Discussion

Estimating cell-type proportions in bulk transcriptomic samples is an important task with a wide spectrum of applications in cancer (i.e., TME estimations can be used to for example predict patient prognosis or response to checkpoint immunotherapy (Sharma, A., et al., 2019; Hendry, S., et al, 2017; Wolf, Y., et al., 2019)). Recently, many digital cytometry tools have attempted to solve this problem mainly by applying or modifying regression approaches (Abbas, A.R., et al., 2009; Newman, A.M., et al., 2015; Wang, X., et al., 2019). Here, we presented ELITE, a new deconvolution method that uses Linear Programming (LP) to solve the blind signal separation (BSS) problem. We showed that ELITE outperforms competing methods in the recovery of known cell fractions when applied to synthetic bulks derived from single-cell RNA-seq data, particularly when the number of cell-types included in the mixtures is high and there are similar cell-types. We also showed the flexibility of ELITE when applied to real bulk samples using publicly available signature matrices that are commonly used, where using a batch-correction scheme not only increases the performance of the deconvolution approach by iteratively transforming the signature matrix, but also generates a signature matrix that biologically aligns better with the bulk samples.

ELITE outperforms competing methods when recovering pseudobulks derived from single-cell data and tumor purity estimations when applied to real bulk data. Its flexibility (not requiring a single-cell dataset as an input to estimate cell fractions) allows ELITE to be applied to publicly available signature matrices (not possible with other algorithms like MuSiC and AdRoit).

When using publicly available single-cell signature matrices, technology differences (microarrays vs sequencing) must be considered. We have coupled ELITE with a batch-correction framework and shown its validity in a large compendium of carcinoma datasets. The correction framework significantly improved the results in 13 of 16 datasets, except for UCEC, LUAD, and LUSC. In UCEC had very comparable results between the raw and batch-corrected solution, however LUAD and LUSC had significantly worse performance. The reason could be that the TR4 matrix was precisely derived from FACS-sorted profiles of epithelial cells (EPCAM+), fibroblasts (CD10+), endothelial cells (CD31+) and immune cells (CD45+), obtained from freshly resected surgical tumor samples from patients with Non-Small Cell Lung Carcinoma (NSCLC) (Newman, A., et al., 2019). Therefore, in these cases we do not need to consider tissue-related differences between TR4, and the mixture matrix M and a different correction approach needs to be used. Future improvements could focus on the automatic identification of these cases. We have also shown that the batch correction produces biologically meaningful signature matrices specific for the tissue type under consideration. We found that it increases the expression of tissue-specific markers while decreasing the expression of lung-specific markers (Table 1).

**Table 1.**
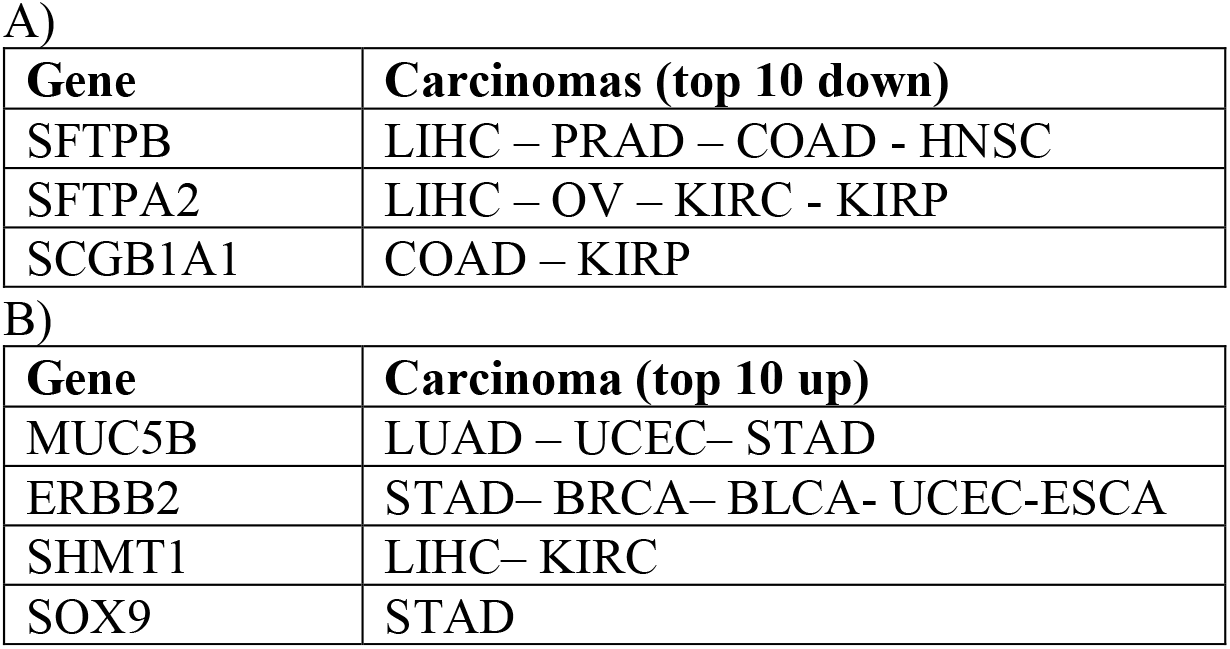
Most variable genes in the carcinoma-specific signature matrices. A) Genes that decreased their value after the optimization process. B) Genes that increased their value after the optimization process. Gene: gene identifier in HGNC; Carcinoma: dataset in which the variation was considered; Positions: For each carcinoma, the corresponding position after sorting by log fold-change.

| Gene | Carcinomas (top 10 down) |
| --- | --- |
| SFTPB | LIHC – PRAD – COAD - HNSC |
| SFTPA2 | LIHC – OV – KIRC - KIRP |
| SCGB1A1 | COAD – KIRP |

B)
| Gene | Carcinoma (top 10 up) |
| --- | --- |
| MUC5B | LUAD – UCEC– STAD |
| ERBB2 | STAD– BRCA– BLCA- UCEC-ESCA |
| SHMT1 | LIHC– KIRC |
| SOX9 | STAD |

Overall, although huge advances have been made in deconvoluting bulk expression datasets, there is still room for improvement. Until scRNA-Seq technology reaches the coverage and cost of bulk experiments, methods such as ELITE will continue to be of high importance to overcome the manifold biological questions emerging from the heterogeneous nature of human cells.

## Acknowledgements

The authors would like to thank PhD Matthew Trotter, PhD Joseph Szustakowski, and PhD Tao Yang for the fruitful discussions and the useful suggestions and comments.

