## supplementary figures for "ELITE: Expression deconvoLution using lInear optimizaTion in bulk transcriptomics mixturEs"

Supplementary Figure 1

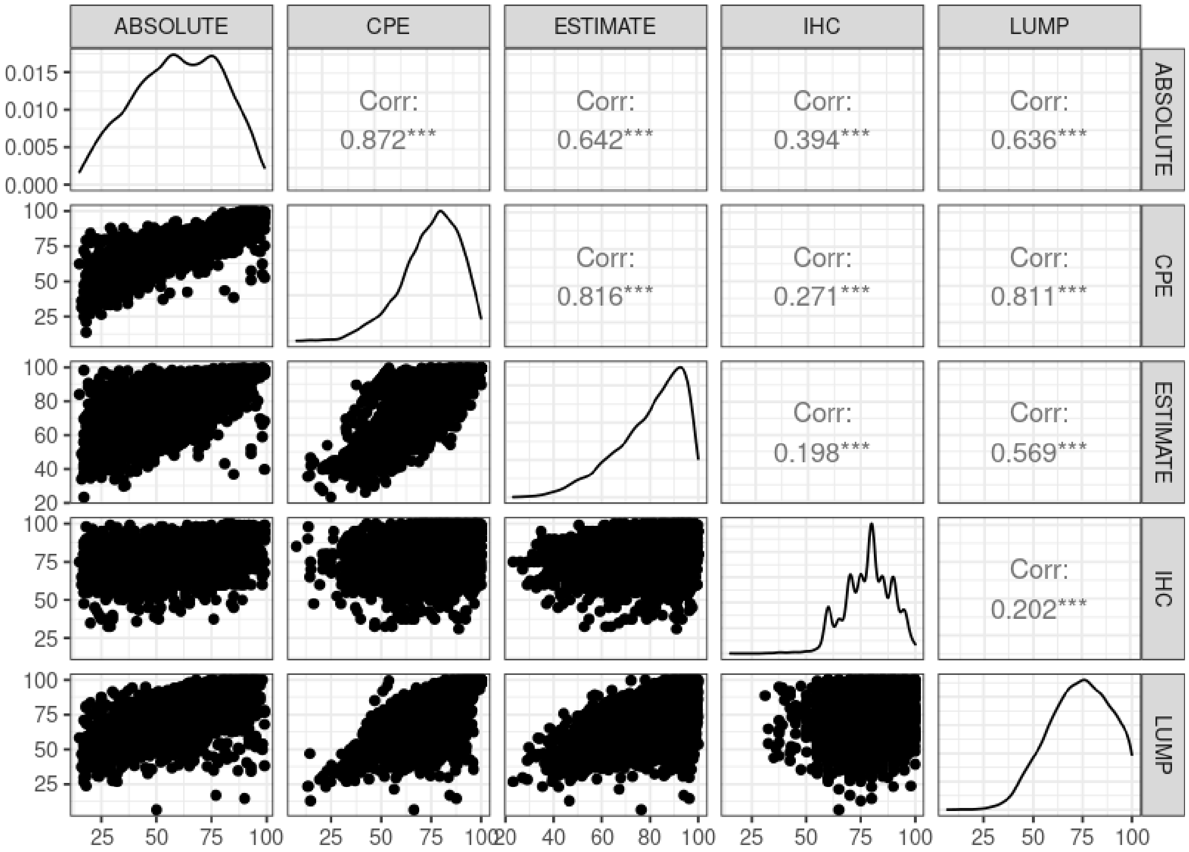


***Supplementary Figure 1 - Correlation between purity estimation methods.*** *Correlation matrix between different purity estimation methods (left to right/top to bottom): ABSOLUTE, CPE, ESTIMATE, IHC, and LUMP. The cells of the lower triangle represent a scatter plot between the purity predictions of the methods described in the specific row/column. The upper triangle shows the Pearson Correlation Coefficient obtained from the comparison of these two methods and the significance associated to the correlation (*** = p<0.001). The diagonal represents a density plot with the distribution of purity estimations for each method. CPE had high PCC with ABSOLUTE (=0.872), ESTIMATE (=0.816), and LUMP (=0.811); ABSOLUTE instead had a moderate PCC with ESTIMATE (=0.642) and LUMP (=0.636); LUMP and ESTIMATE also had a moderate PCC between them (=0.569); and IHC had a low PCC with any of the other methods (=0.394, =0.271, =0.198, and =0.202 for ABSOLUTE, CPE, ESTIMATE, and LUMP respectively).*

Supplementary Figure 2

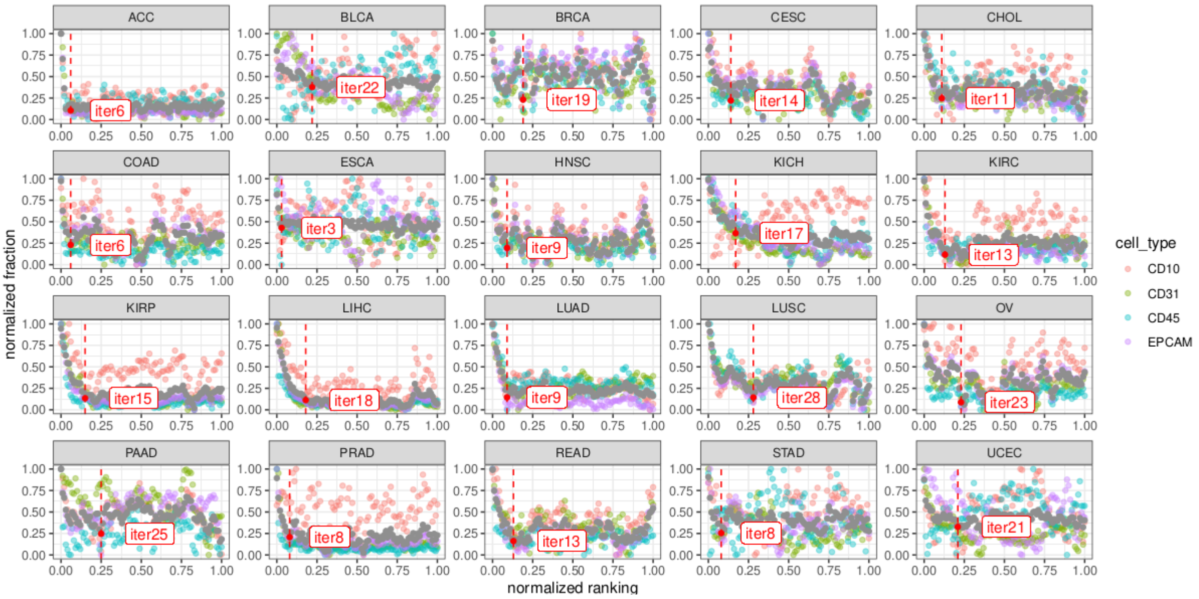


*Supplementary Figure 2 - Selection of the optimal iteration. Scatter plot showing the normalized fraction (y-axis) for each iteration (normalized ranking, x-axis). The gray dots represent the average at each iteration. The elbow criterion was used as stopping criteria (i.e., closest point to the (0,0) origin).*

Supplementary Figure 3

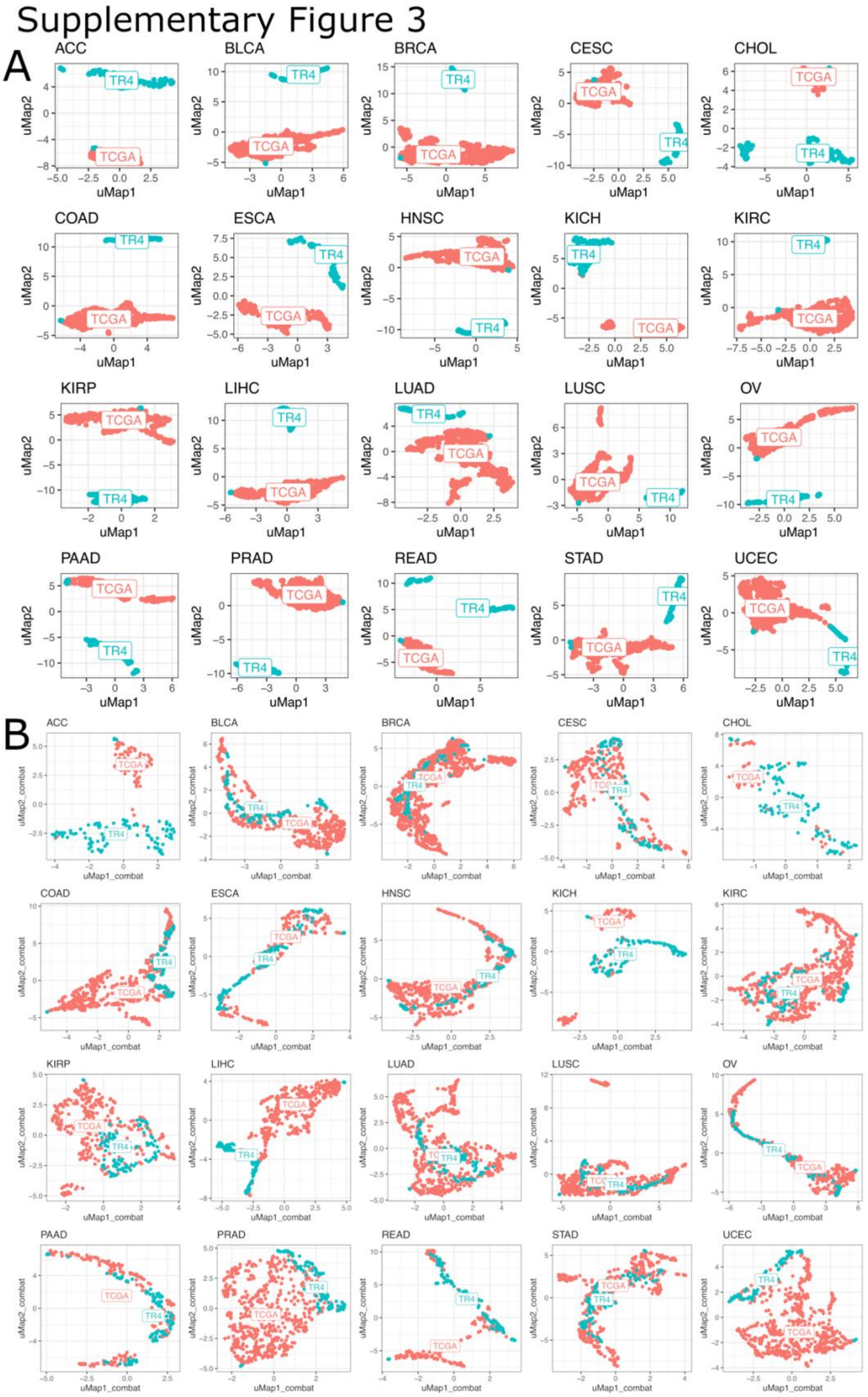


*Supplementary Figure 3 - Batch-correction using ComBat-seq. Scatter plots showing the umap before (A) and after (B) correction with ComBat-seq. Each subplot represents a different TCGA carcinoma dataset. Red dots represent samples while blue dots represent columns of TR4 or pseudobulks generated after a weighted sum of the columns of TR4.*

Supplementary Figure 4

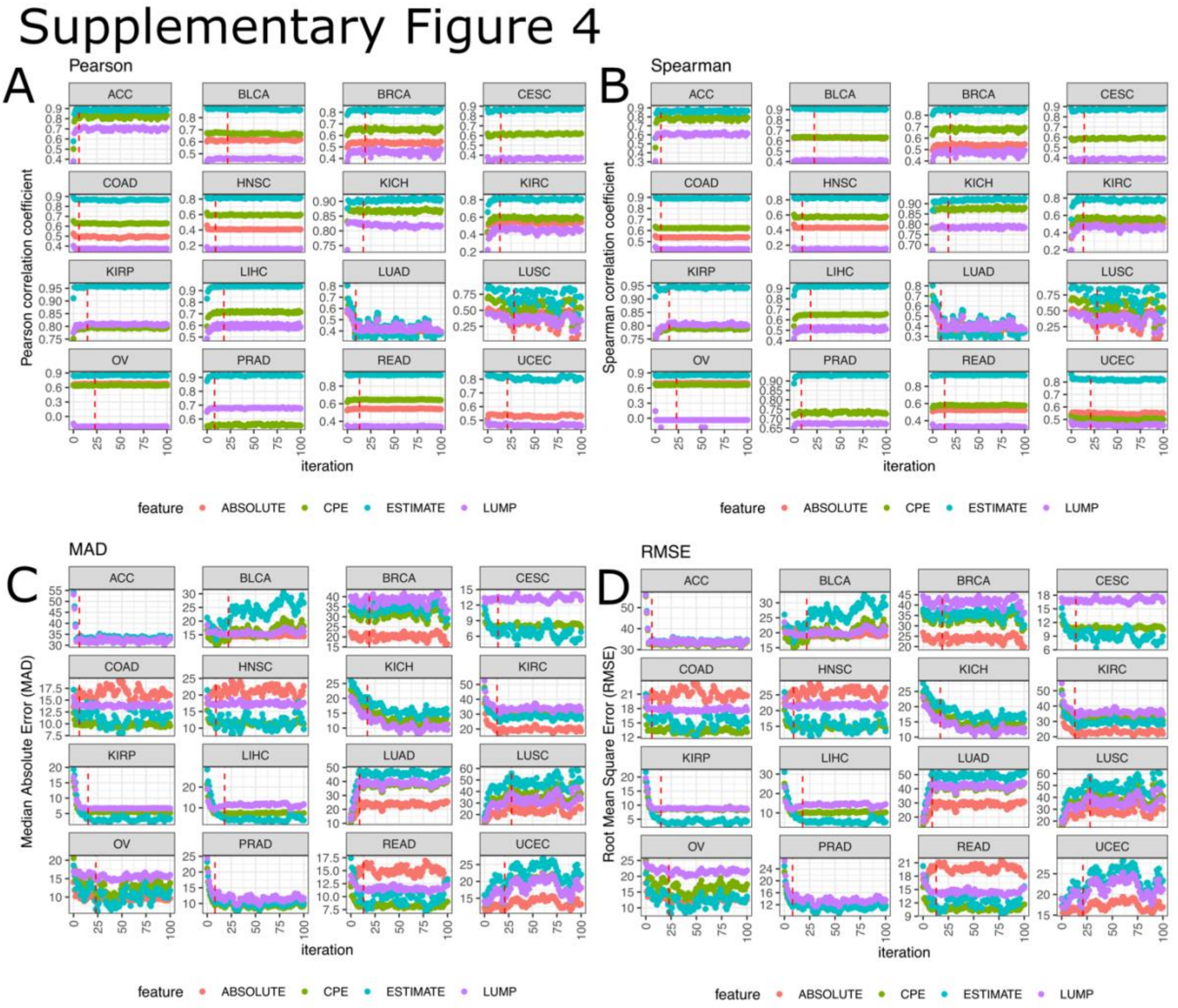


*Supplementary Figure 4 - Evolution graphs for the performance of the batch correction iterations. Scatter plots showing the Pearson Correlation Coefficient (PCC), Spearman Correlation Coefficient (SCC), Median Absolute Error (MAD), and Root-Mean Square Error (RMSE) for each dataset, comparing the predicted fraction of tumoral cells by ELITE in each iteration (x-axis) and the purity estimation using the different purity estimation methods (ABSOLUTE, CPE, ESTIMATE, and LUMP).*

Supplementary Figure 5

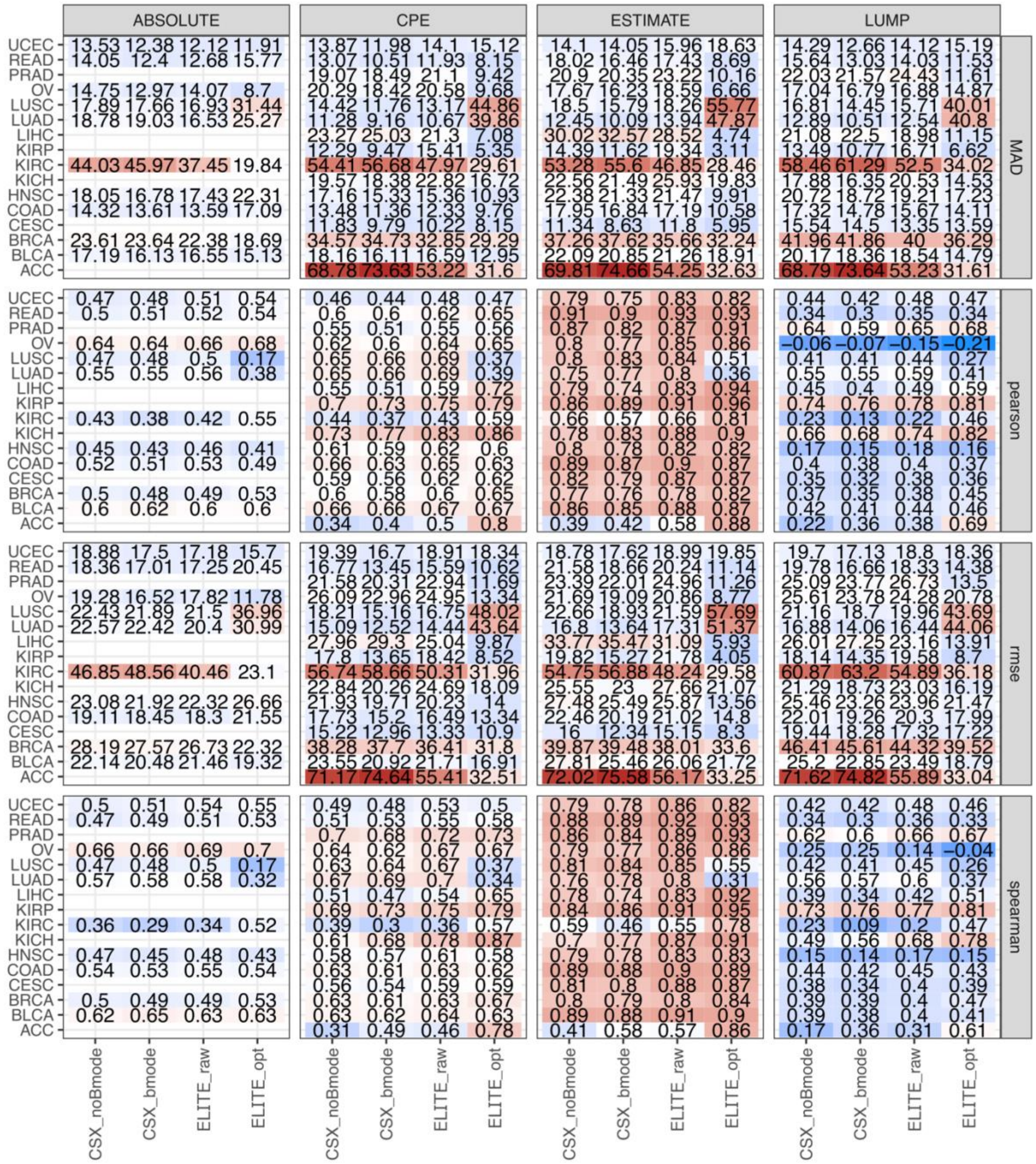


***Supplementary Figure 5 - Comparison of different deconvolution method in real bulk samples.*** *Heatmap showing the performance of 20 different carcinoma datasets from TCGA using four different implementations (CSX without batch effect correction, CSX with batch effect correction, ELITE without batch effect correction, and ELITE with batch effect correction). The heatmaps include 4 different statistics (Pearson Correlation Coefficient, Spearman Correlation Coefficient, Median Absolute Error, and Root-Mean-Square Error) in four different tumor purity estimation methods (ESTIMATE, ABSOLUTE, LUMP, and CPE).*

*Supplementary Table 1 – TCGA carcinoma datasets used for the validation of bulk RNA-seq data.*

| Study Abbreviation | Study Name | N |
|---|---|---|
| ACC | Adrenocortical carcinoma | 79 |
| BLCA | Bladder Urothelial Carcinoma | 430 |
| BRCA | Breast invasive carcinoma | 1,217 |
| CESC | Cervical squamous cell carcinoma and endocervical adenocarcinoma | 309 |
| CHOL | Cholangiocarcinoma | 45 |
| COAD | Colon adenocarcinoma | 512 |
| ESCA | Esophageal carcinoma | 173 |
| HNSC | Head and Neck squamous cell carcinoma | 546 |
| KICH | Kidney Chromophobe | 89 |
| KIRC | Kidney renal clear cell carcinoma | 607 |
| KIRP | Kidney renal papillary cell carcinoma | 321 |
| LIHC | Liver hepatocellular carcinoma | 424 |
| LUAD | Lung adenocarcinoma | 585 |
| LUSC | Lung squamous cell carcinoma | 550 |
| OV | Ovarian serous cystadenocarcinoma | 379 |
| PAAD | Pancreatic adenocarcinoma | 182 |
| PRAD | Prostate adenocarcinoma | 551 |
| READ | Rectum adenocarcinoma | 177 |
| STAD | Stomach adenocarcinoma | 407 |
| UCEC | Uterine Corpus Endometrial Carcinoma | 583 |
